# Structural determinants for GqPCR-mediated inhibition of TASK K2P K^+^ channels and their dysfunction in disease

**DOI:** 10.1101/2025.08.04.668454

**Authors:** Thibault R. H. Jouen-Tachoire, Peter Proks, David Seiferth, Kate Crowther, Philip C. Biggin, Thomas Baukrowitz, Marcus Schewe, Stephen J. Tucker

## Abstract

Two-Pore Domain K^+^(K2P) channels are crucial determinants of cellular electrical excitability. TASK-1 and TASK-3 K2P channels regulate the resting membrane potential in many different cell types where their activity is also coupled to GqPCR signalling pathways via direct inhibition by diacylglycerol (DAG) generated as a result of phosphatidylinositol-4,5-bisphosphate (PIP_2_) hydrolysis. This regulation is defective in two different TASK channelopathies, but the molecular mechanisms underlying this inhibition remain unclear. Here, we demonstrate that DAG inhibition of TASK channel activity is state-dependent. Single channel recordings show that the sensitivity to GqPCR inhibition inversely correlates with channel open probability and that DAG destabilises the open state of TASK-1 to promote channel closure. Combining Molecular Dynamics simulations with mutagenesis studies, we also identify a binding site for DAG in a groove between the M2, M3 and M4 domains, and highlight the crucial role of a specific residue within on M4 (T230) in mediating this inhibitory effect as well as defining the difference in GPCR sensitivity between TASK-1 and TASK-3. Together, these results provide a better understanding of the molecular mechanisms underlying GqPCR regulation of TASK channels and the pathogenic effect of K2P channelopathies linked to TASK channel dysregulation.

## Introduction

Potassium (K^+^) channels are transmembrane pores that allow the selective diffusion of K^+^across cellular membranes to control cellular electrical activity. In particular, the family of Two-Pore Domain K^+^ (K2P) channels are important for the generation and regulation of the resting membrane potential (*1*). However, although they were originally described as ‘leak’ channels, they are now known to be regulated by diverse physico-chemical stimuli, as well as a range of GPCR-dependent and other cellular and metabolic signalling pathways (*2*).

There are 15 different human K2P channels encoded by the *KCNK1-18* genes and are grouped according to both their sequence similarity and functional properties. In particular, TASK K2P channels include two members, TASK-1 (*KCNK3*) and TASK-3 (*KCNK9*) which were originally characterised by their sensitivity to inhibition by external protons (H^+^) across the physiological pH range, a property from which they derive their name (<u>T</u>WIK-related <u>A</u>cid <u>S</u>ensitive <u>K</u>^+^ channels) (*3*). GPCR-coupled signalling has also been shown to inhibit TASK channels, and provides a pathway for neurotransmitters and hormones to influence cellular excitability (*4*). This GPCR-modulation of TASK currents also plays an important role in the control of aldosterone secretion and a variety of pathways involved in the mechanical control of ventilation as well as both central and peripheral chemosensation (*5–7*). TASK channels are also found in certain nociceptive sensory neurons, as well as the heart, and their crucial role in regulating electrical activity in these various tissues means they have also been proposed as targets for the treatment of pain, atrial fibrillation and sleep apnea (*8–10*).

The GPCR-mediated inhibition of TASK channels has been proposed to involve many different factors (*4*) though is principally via Gαq-coupled receptors (*11–13*). Interestingly, an elegant dissection of this pathway demonstrated a critical role for activation of Phospholipase C (PLC) which triggers hydrolysis of PIP_2_ into soluble inositol-1,4,5-trisphosphate (IP_3_) and membrane-bound diacylglycerol (DAG). PIP_2_ is a known activator of many different K^+^ channels, but it is not the depletion of PIP_2_ *per se* which leads to a loss of channel activity, instead it is the consequent increase in DAG which leads to a direct inhibition of TASK currents (*14*). However, the biophysical, kinetic and molecular mechanisms that underlie this direct interaction and ultimate modulation of channel gating remain unknown. TASK channels lack the classical cysteine rich C1 domains that typically bind DAG, and DAG lacks the more complex headgroup and negative charges that define many phospholipid/channel interactions, especially as seen with PIP_2_.

Recently we identified a novel TASK-1 channelopathy, Developmental Delay with Sleep Apnea (DDSA) associated with a range of *de novo* gain-of-function (GoF) heterozygous missense mutations in *KCNK3 (15)*. The mutations cluster near the X-gate, a gating motif which regulates intracellular access to the inner cavity and which has previously been implicated in channel activation by volatile anaesthetics, as well as GqPCR regulation (*16, 17*). We demonstrated that these DDSA mutations increase channel open probability (*P*_*o*_) and that their GoF effect is exacerbated by a markedly reduced sensitivity to GqPCR inhibition due to a concomitant reduction in sensitivity to inhibition by DAG. Interestingly, similar functional effects are also seen with a number of GoF mutations in TASK-3 associated with another neurodevelopmental disorder, *KCNK9* Imprinting syndrome (KIS) (*18, 19*). This therefore suggests that there may be a common gating mechanism that is defective in all these disease-causing mutations in TASK channels which may provide insight into the molecular basis of their regulation by DAG and GPCR-dependent regulation. Recent structural studies of TASK channels also now provide a wealth of structural information to support such analysis (*20–22*).

In this study, we have used both macroscopic and single channel recordings of TASK channels to examine their GPCR-mediated inhibition and demonstrate a state-dependent mechanism of regulation by DAG. We also examine the molecular determinants of this DAG-TASK channel interaction and identify a specific site that determines the difference in GPCR sensitivity between TASK-1 and TASK-3. These results not only advance our understanding of the molecular mechanisms underlying GPCR regulation of TASK channels but also the pathogenic effect of TASK channelopathies linked to channel dysregulation.

## Methods

### Molecular biology

The wild-type human TASK-1 gene (*KCNK3*) and human TASK-3 gene (*KCNK9*) in the pFAW dual purpose oocyte expression vector were used throughout this study (*23*). For GqPCR regulation, channels were coexpressed with the human P2Y2 receptor (*15*). Mutations in TASK channels were introduced by site-directed mutagenesis and confirmed by sequencing. RNA was made using the T7 mScript Standard mRNA Production System for expression in *Xenopus* oocytes.

### Electrophysiology

*Whole-cell recordings:* each oocyte was injected with 2 ng of RNA and incubated for 20-24 h at 17.5° C in Modified Barth’s storage solution at pH 7.4 (88 mM NaCl, 1 mM KCl, 1.68 mM, MgSO_4_7H_2_O, 10 mM HEPES, 2.4 mM NaHCO_3_) supplemented with 0.05 mg/ml gentamicin. Recordings were performed in ND96 buffer at pH 7.4 (96 mM NaCl, 2 mM KCl, 2 mM MgCl_2_, 1.8 mM CaCl_2_, 5 mM HEPES). Currents were recorded using a 400 ms voltage step protocol from a holding potential of −80 mV delivered in 10 mV increments between −120 mV and +50 mV followed by 800 ms ramp protocols from −120 to +50 mV. GqPCR inhibition studies were conducted as before (*15*) where currents were recorded before and after perfusion with 15 µM ATP, a concentration that elicits 50% inhibition of WT TASK-1. All recorded traces were analyzed using Clampfit (Axon Instruments), and graphs were plotted using Origin2021 (OriginLab Corporation).

### Single channel recordings

Single channel currents were recorded with Axopatch 200B amplifier via a Digidata 1440A digitizer (Molecular Devices) at −100 mV where the channels have sufficiently large single-channel conductance suitable for analysis of their gating. Data were filtered at 2 kHz and recorded at a 200 kHz sampling rate with program Clampex (Molecular Devices). Pipette solution and bath solution for cell-attached experiments contained (in mM): 140 KCl, 2 MgCl_2_, 1 CaCl_2_, 10 HEPES (pH 7.4 adjusted with KOH/HCl). For inside-out experiments, the bath solution contained (in mM): 140 KCl, 2 MgCl_2_, 1 CaCl_2_, 10 HEPES (pH 7.2 adjusted with KOH/HCl). All experiments were conducted at room temperature. For analysis of single-channels, open probability was calculated from idealized current traces constructed with 50 % threshold criterion with Clampfit (Molecular Devices) at an imposed resolution of 50 μs. Current data points below threshold (half of the open level) were interpreted as closed state and data points above threshold as open state. Horizontal lines going through means of the current values of individual openings and closings were generated using Clampfit. Summing the lengths of all openings and closings in the idealized recording allowed the program to calculate *P*_*o*_. Analysis of amplitude and dwell-time distributions was performed in Origin (Origin-Lab Corporation). The critical time for burst analysis was determined using Magleby and Pallotta criterion (*24*). Statistical significance in DiC8 effects on channel lifetimes was determined by paired t-test. Dwell-time histograms are presented using the Sine-Sigworth display, where the x-axis is the logarithm of a state duration and the y-axis is its corresponding square root number of occurrences in the recording. The apparent kinetic states of the channel manifest themselves as peaks in the histogram with their mean lifetimes being the maximum peak values on the y-axis. Bursts are defined as composite states consisting of openings and short closings The lines are best fit of probability density functions *pdf(t)=Σa*_*i\**_*exp(−t/t*_*i*_) to the data where *t*_*i*_ is a mean lifetime of a state *i* and *a*_*i*_ is the corresponding relative area of events of the state *i* in the distribution (*Σa*_i_=1).

### Macroscopic patch recordings

Giant excised membrane patch measurements in inside-out configuration under voltage-clamp conditions were made at room temperature 72-120 h after injection of 50nl channel specific mRNA into oocytes. Thick-walled borosilicate glass pipettes had resistances of 0.25 - 0.35 MΩ (tip diameter of ~15 - 30 µm) and were back-filled with extracellular solution containing (in mM): 4 KCl, 116 NMDG, 10 HEPES and 3.6 CaCl_2_ (pH was adjusted to 7.4 with KOH/HCl). Bath solution was applied to the cytoplasmic side of the excised giant patches via a gravity flow multi-barrel application system and had the following composition (in mM): 120 KCl, 10 HEPES, 2 EGTA and 1 Pyrophosphate. Currents were acquired with an EPC10 USB amplifier and HEKA PatchMaster 2×91 software. The sampling rate was 10 kHz and the analog filter was set to 3 kHz (−3 dB). Voltage ramp pulses (−80 mV to +80 mV) were applied from a holding potential (V_H_) of −80 mV for 0.8 s with an inter-pulse interval of 9 s and were analyzed at a given voltage of +40 mV. The relative steady-state levels of inhibition for indicated blocker were fitted with the following Hill equation:

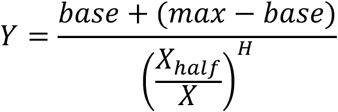

where *base* is the inhibited (zero) current, *max* is maximum current, *X* is the blocker concentration, *X*_*half*_ is the value of concentration for half-maximal occupancy of the blocker binding site and *H* is the Hill coefficient. 1,2-Dioctanoyl-*sn*-glycerol (DiC8) were purchased from Sigma-Aldrich. All compounds were stored as stock solutions (10-100 mM) at −80° C and were diluted in intracellular bath solution to final concentrations prior to each measurement.

### Molecular dynamics

In the coarse-grained simulations, three 20 μs simulation repeats were performed using the Martini 2.2 forcefield with an elastic network (EINeDyn) (*25*). POPC membranes with different concentrations (5 %, 10 %, 20 %, 33 %) of DAG in the lower leaflet were tested. Binding sites were characterized by residues with the highest residence time and occupancy, employing a dual-distance cutoff approach with pyLipID which uses a community analysis approach for binding site detection, calculating lipid residence times for the individual protein residues and the detected binding sites (*26*). For representative binding poses generated with coarse grained simulations, the cg2at.py tool was used to map coarse-grained beads to atoms and evaluated the stability of the binding poses with atomistic simulations (*27*). Atomistic simulations were then set-up from scratch using Charmm GUI, with DAG concentrations (20 %) in the lower leaflet and varying sidechain lengths and saturation (C12:0/C12:0, C14:0/C14:0, C16:0/16:0, C16:1/16:1, and C16:0/18:2). Simulations were performed with GROMACS 2021, using the TIP3P water model and the CHARMM36m forcefield (*28*). pyLipID was used to assess the DAG and protein interactions in three independent 5 µs long atomistic repeats.

## Results

### Mutant TASK channels with a range of intermediate open probabilities

We previously reported that all of the DDSA mutant channels identified so far had a markedly reduced GqPCR sensitivity with relatively little difference between them, i.e., there were none with an ‘intermediate’ or only minor reduction in GqPCR sensitivity (*15*). Likewise, they all exhibited a marked increase in single channel *P*_*o*_ of at least 10-fold. One possible mechanism that could explain this correlation is a state-dependent mechanism of action of DAG on TASK channels whereby the ligand interacts preferentially with either the closed or open state to exert its functional effect. However, to examine this more fully requires the identification of additional mutant channels with a wider range of *P*_*o*_ to enable comparison with their GqPCR inhibition and sensitivity to DAG. Our initial screen therefore focussed on introducing a variety of different amino acid substitutions at three of the key sites mutated in DDSA (F125S, Q126E and G129S) that produce channels with a relatively high *P*_*o*_ and markedly reduced GqPCR sensitivity (*15, 29*).

Initial two-electrode voltage-clamp (TEVC) recordings show that macroscopic, whole-cell current levels for many substitutions at these positions exhibited an increase in whole-cell activity to a range of different levels (**Figure 1a**). We therefore next examined their single channel behaviour and found that they also exhibited a suitable range of open probabilities (**Supplementary Table S1**) such as those seen with different substitutions at G129 (**Figure 1b**).

**Figure 1.**
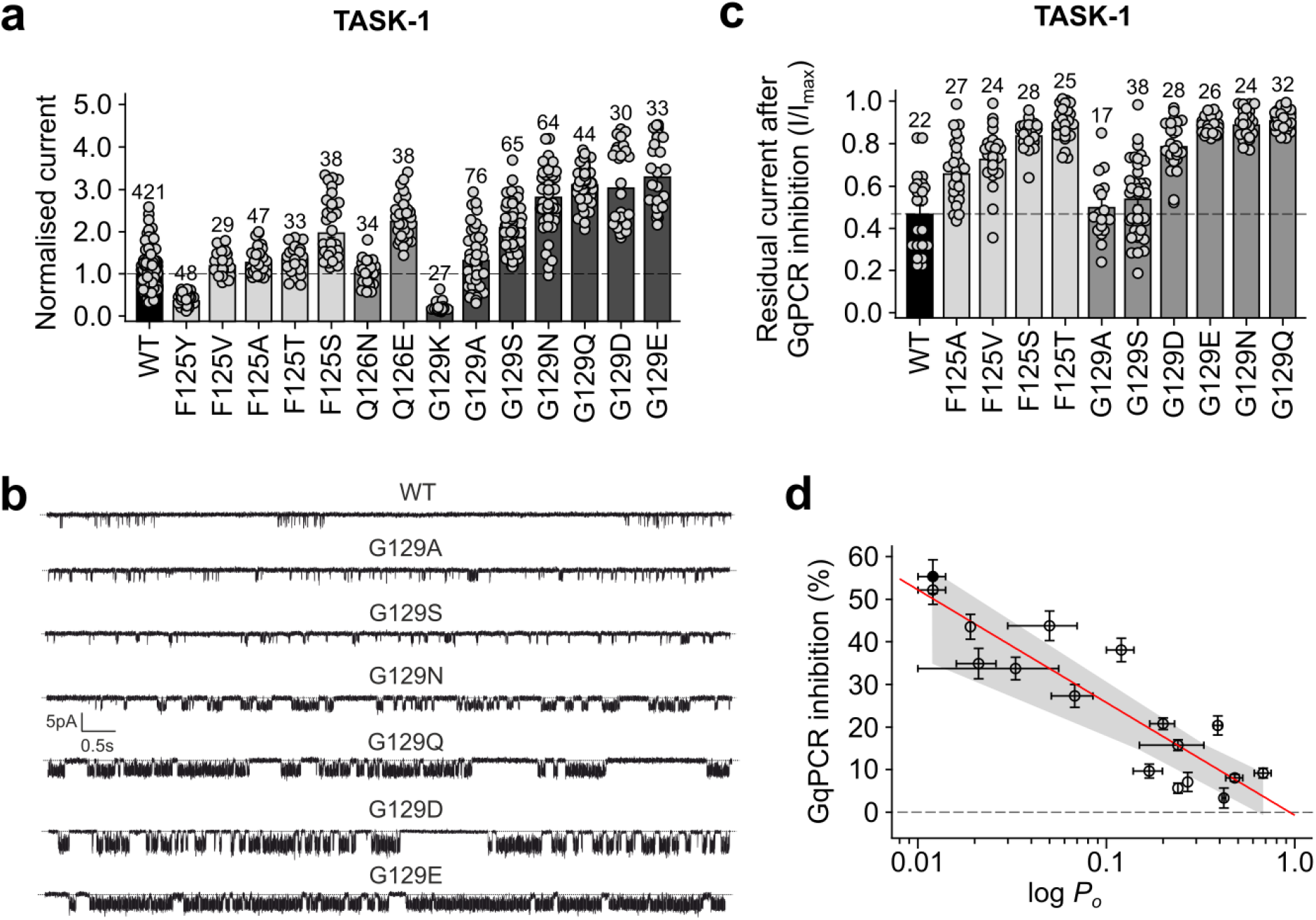
Correlation between TASK-1 GqPCR sensitivity and channel *P*_*o*_. **a**. Macroscopic whole-cell currents for different mutations at positions F125, Q126 and G129. b. Representative single channel recordings of WT and mutant TASK-1 channels in cell-attached patches recorded at −100mV showing a range of *P*_*o*_ **c**. Relative GqPCR inhibition for GoF variants shown in panel a. **d**. Steep inverse correlation between GqPCR inhibition and single channel *P*_*o*_ (y = −26.49; R^2^ = −0.79) with the linear regression shown as red line and the 95 % confidence intervals of the fit shown as the shaded grey area. WT TASK-1 is shown as solid black circle.

### An inverse correlation between GqPCR sensitivity and P_o_

We next measured mutant channel sensitivity to GqPCR inhibition via coexpression with the P2Y2 receptor (*15, 30, 31*). Importantly, we found that those mutations which exhibited a single channel *P*_*o*_ between 0.01 and 0.7 also exhibited a corresponding range of sensitivities to GqPCR-mediated inhibition (**Figure 1c**). To better understand this relationship, we plotted these *P*_*o*_ values and observed a clear correlation where GqPCR sensitivity decreases sharply as channel *P*_*o*_ increases. This inverse relationship is most clearly visualised in the plot shown in **Figure 1d**.

### Dynamic modulation of channel activity alters GqPCR sensitivity

The mutations examined above have their limitations due to the varying effects that some mutations can have on the properties of ion channels, not just their overall *P*_*o*_. A far better method would therefore be to dynamically modulate the activity of WT TASK channels and measure this effect on GqPCR sensitivity. Interestingly, both TASK-1 and TASK-3 have been shown to be activated by volatile anaesthetics and a study of TASK-3 previously reported that its GqPCR sensitivity is reduced upon activation by halothane (*16*) or chemical modification (*32*). We therefore examined the effect of other mechanisms of channel activation and inhibition on GqPCR sensitivity without the need for mutagenesis.

TASK-1 is known to be inhibited by extracellular H^+^ within the physiological range and so we first examined the effect of changing extracellular pH (pH_e_). The *pK*_*a*_ of TASK-1 is 7.4 so to avoid any possible effect of changes in pH on the ATP ligand itself (*pK*_*a*_ 6.4), we compared GqPCR inhibition at pH 8.5 compared to pH 7.4. Interestingly, we found that the extent of GqPCR inhibition was indeed reduced at pH_e_ 8.5 where TASK-1 channels are activated compared to pH_e_ 7.4 (**Figure 2a**).

**Figure 2.**
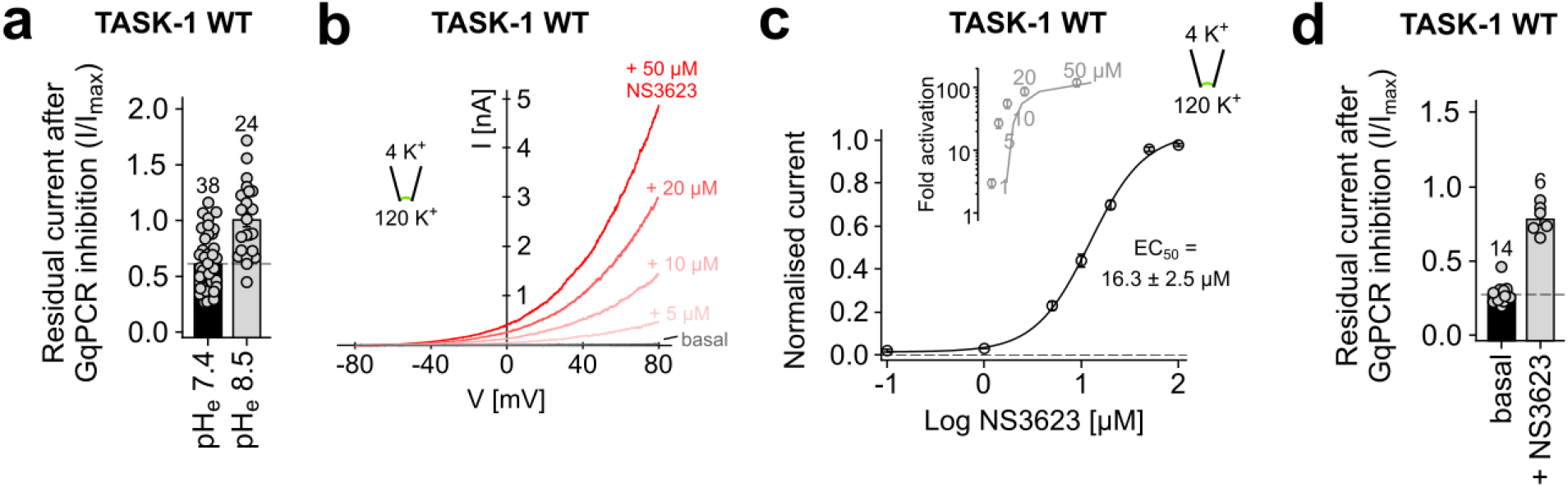
Dynamic modulation of channel activity by pH_e_ and NS3623 alters GqPCR-sensitivity. **a**. Residual whole-cell current after GqPCR inhibition recorded at pH_e_ 7.4 vs pH_e_ 8.5 where TASK-1 activity is higher (*P* < 0.001) and GqPCR inhibition is reduced. **b**. Activatory effect of different concentrations of NS3623 on WT TASK-1 currents recorded in giant excised patches. **c**. Dose-response curve for TASK-1 activation by NS3623. Inset (grey) is shown the fold-activatory effect of indicated concentrations. **d**. Whole-cell WT TASK-1 currents activated by NS3623 (50 µM) exhibit a markedly reduced GqPCR inhibition (*P* < 0.001).

No small molecule drugs have currently been shown to directly activate TASK-1 channels via an increase in *P*_*o*_. A class of negatively-charged activators (NCAs) directly activate many different K_2P_ channels, but the most common, BL-1249, has little effect on TASK channels (*33*). We therefore examined whether other ligands with a similar chemical structure to the NCA pharmacophore could directly activate TASK-1. We found that a related compound, NS3623 that has previously been shown to activate Kv4.3 and hERG K^+^ channels (*34*), potently activated TASK-1 in excised patches (**Figure 2b-c**). Single channel recordings in cell-attached configuration at −100mV also showed that this activation directly increased channel *P*_*o*_ from 0.022 ± 0.003 (*n* = 10) to 0.49 ± 0.06 (*n* = 10).

We therefore measured whether activation by NS3623 also shifted GqPCR sensitivity. Consistent with our findings, we found that NS3623 markedly reduced the extent of GqPCR-mediated inhibition (**Figure 2d**). Overall, these results are therefore consistent with a state-dependent mechanism for GqPCR-inhibition of TASK-1.

### GqPCR inhibition correlates with changes in sensitivity to inhibition by DAG

GqPCR-mediated inhibition of TASK channels is via direct interaction of DAG and we have previously shown that the DDSA mutation, N133S which has a reduced GqPCR sensitivity exhibits a correspondingly dramatic reduction in sensitivity to inhibition by a short acyl chain derivative of DAG, *i*.*e*., 1,2-dioctanoyl-sn-glycerol (DiC8) when applied directly to excised patches from the intracellular side (*15*). If the effect of this interaction is state-dependent, then we would expect those mutations with a range of single channel *P*_*o*_ to exhibit corresponding reductions in their sensitivity to DAG.

We therefore examined DiC8 inhibition on different substitutions at G129 which exhibit a suitable range of *P*_*o*_ and GPCR sensitivities. This revealed a range of reduced sensitivity to DiC8 and IC_50_ values that correlated with their relative increase in *P*_*o*_ (**Figure 3a-c**). We next investigated whether dynamic modulation of WT TASK-1 activity also affected DiC8 inhibition in excised patches and found that NS3623 activation markedly reduced this inhibitory effect (**Figure 3d-e**).

**Figure 3.**
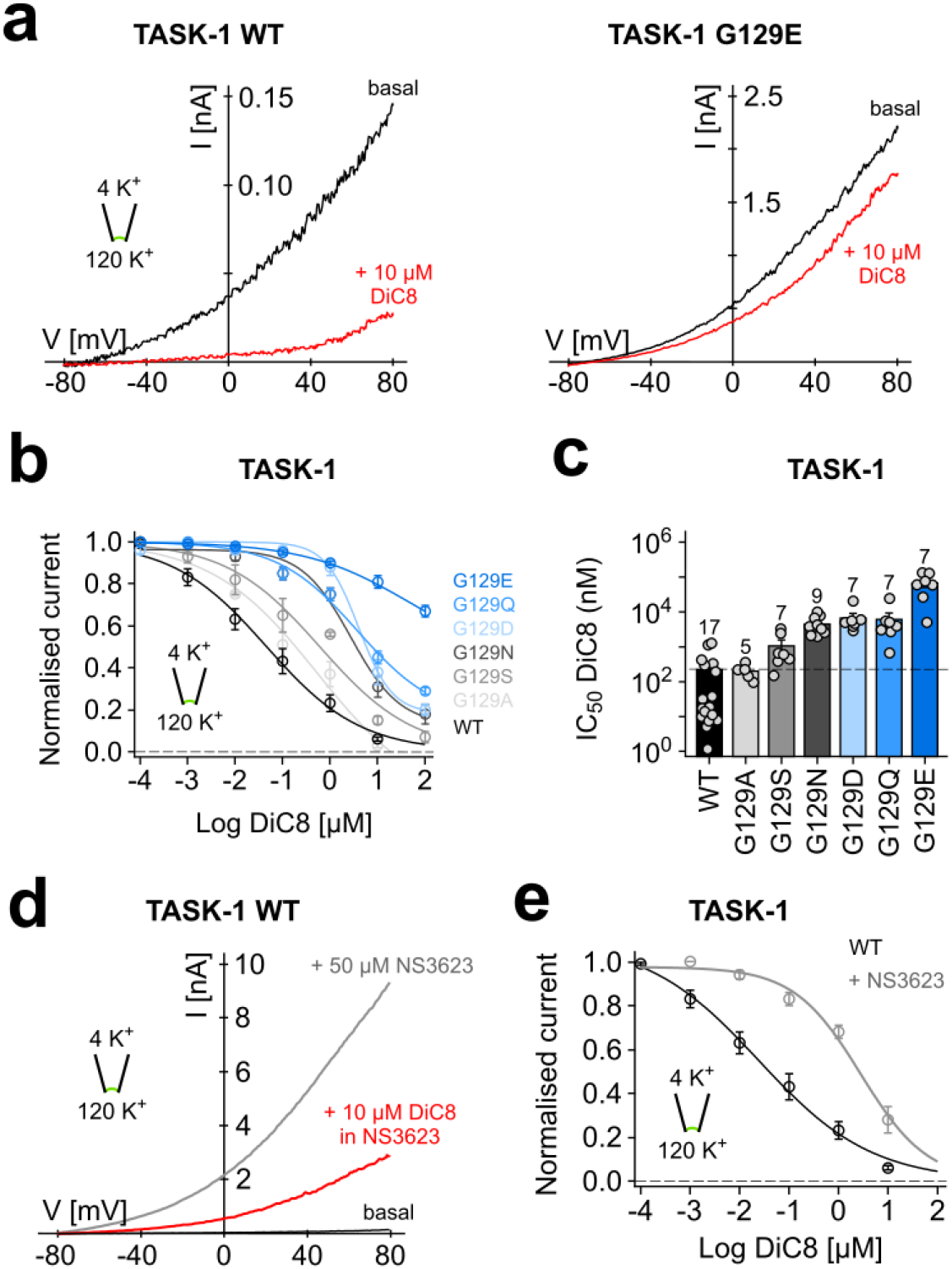
Direct inhibition of WT and mutant TASK-1 channels by the synthetic DAG homologue DiC8. **a**. Inhibition of WT TASK-1 vs. TASK-1 G129E by a short chain DAG analogue (10 µM DiC8). Currents recorded in giant excised patches and DiC8 applied to the intracellular side of the respective patch. **b**. Dose-response curves calculated for inhibition of WT and mutant TASK-1 showing a variety of responses to DiC8 inhibition. **c**. IC_50_ values for G129 mutants calculated from data shown in panel b. **d**. Activation of WT TASK-1 by NS3623 markedly reduces the inhibitory effect of 10 µM DiC8. **e**. Dose-response curve showing the marked reduction in DiC8 inhibition after activation of WT TASK-1 with 50 µM NS3623.

Overall, these effects strongly support a mechanism for state-dependent inhibition of TASK channels via a direct interaction with DAG. However, such recordings of macroscopic currents cannot distinguish between the open and closed states of TASK channels. Therefore, to further evaluate this mechanism we examined the inhibitory effect of DiC8 on TASK-1 at the single channel level.

### DAG destabilises the open state of TASK-1

If DAG inhibits TASK channel activity via a reduction in *P*_*o*_, then this can occur in at least two ways; either via stabilisation of the closed states of the channel, or by destabilising the open states because the resulting macroscopic effect will be the same. However, TASK-1 has a very low intrinsic channel *P*_*o*_ and so to measure the effects of DAG inhibition on single WT channels would be challenging. We therefore examined the effect of DiC8 on TASK-1 N133S mutant channels in excised patches because this mutant has a *P*_*o*_ ~0.2 compared to ~0.01 for WT TASK-1, but can still be completely inhibited by DiC8, albeit at much higher concentrations than for WT TASK-1 (*15*). This showed that application of 1 µM DiC8 produced a dramatic reduction in single channel *P*_*o*_. (**Figure 4a**)

**Figure 4.**
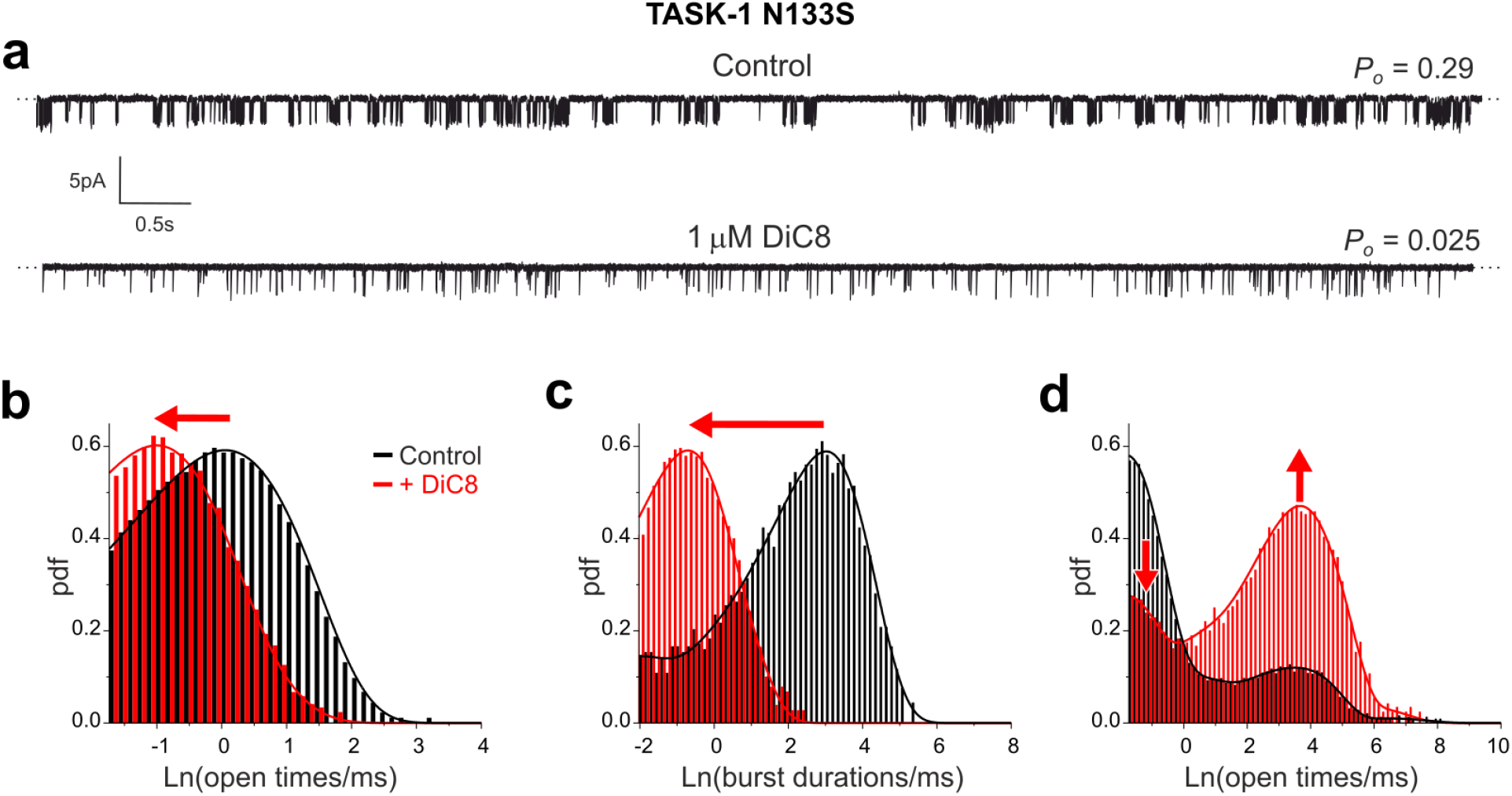
DiC8 destabilises the open state of TASK-1. **a**. Single channel traces showing inhibition of N133S TASK-1 channels by 1µM DiC8 applied to the inside of an excised patch (bottom trace). The sample traces are 1 s long measured at −100mV. **b-d**. Dwell-time histograms (presented using Sine-Sigworth display) are obtained from the full recordings (> 4 minutes) either in the absence (black) or presence (red) of DiC8 and are overlayed for comparison. The apparent kinetic states of the channel manifest themselves as peaks in the histogram with their mean lifetimes being the maximum peak values. Three different types of distributions are shown as indicated: (**b**) distribution of openings, (**c**) bursts of opening, and (**d**) closings. The bursts are composite states consisting of openings and short closings contained within the peak on the far left in the distribution of closings. The lines are the best fit of probability density functions (see methods for details and also **Supplementary Table S2**). The red arrows indicate the direction of shift in the major peaks during inhibition by DiC8.

Comparison of the dwell-time histograms for TASK-1 N133S before and after application of DiC8 reveal that it substantially reduced the duration of openings and burst durations (**Figure 4b-c**). The distribution of closed times consists of two major peaks which correspond to the intra-burst and inter-burst closed states, and a small population of very long closed states (**Figure 4d**). DiC8 did not appear to noticeably alter the relative position of these two major peaks; however, it markedly reduced the number of short (intra-burst) closed states and concomitantly increased the number of long (inter-burst) closed states (**Figure 4d**).

This suggests that open state destabilisation by DiC8 leads to more frequent exits from the open states to inter-burst closed states which dramatically increases their number (**Figure 4d**). This is not the case for the short closed states within bursts, thus their relative number in the presence of DiC8 are diminished. This effect also contributes to the overall reduction in burst durations (**Figure 4b**). A more precise analysis of the intra-burst closures is limited by the temporal resolution of the acquired data; however, their potential destabilisation would further reduce both their relative number as well as the length of bursts. Overall, these results therefore imply that DiC8 inhibits TASK channels primarily via destabilisation of the open states and bursts of channel openings.

### DAG binds to TASK-1 in a groove between M2, M3 and M4

Our results support a mechanism where direct binding of DAG to TASK-1 reduces channel *P*_*o*_, but the precise site of interaction between DAG and TASK-1 remains unknown. High-resolution structural information now exists for both TASK-1 and TASK-3 and so we chose to take advantage of recent advances in the analysis of molecular dynamics (MD) simulation of lipid/ protein interactions in membranes to probe for potential DAG binding sites.

PyLipID is a Python package that uses a community analysis approach for lipid binding site detection and calculating lipid residence times for both the individual protein residues and the detected binding sites (*26*). Coarse-grained simulations were therefore initially performed with different numbers of DAG molecules in the lower leaflet. Potential interactions were then characterised by using PyLipID to identify residues with the highest residence time and occupancy. Representative binding poses were then converted into full-atom models for further atomistic simulations with DAG in the lower leaflet. A similar PyLipID analysis of these simulations revealed that the headgroup of DAG exhibits the highest residence time and occupancy with two main residues on M4, T230 and V234, both within a membrane-facing groove between the M2, M3 and M4 transmembrane helices of same subunit of the channel (**Figure 5a**). Varying the length and degree of saturation of the DAG acyl chains did not have any major influence on interaction with this site, though consistent with the mobility of DAG within the membrane (*35*), the acyl chains were capable of adopting several poses within this groove (**Supplementary Figure S1a)**.

**Figure 5.**
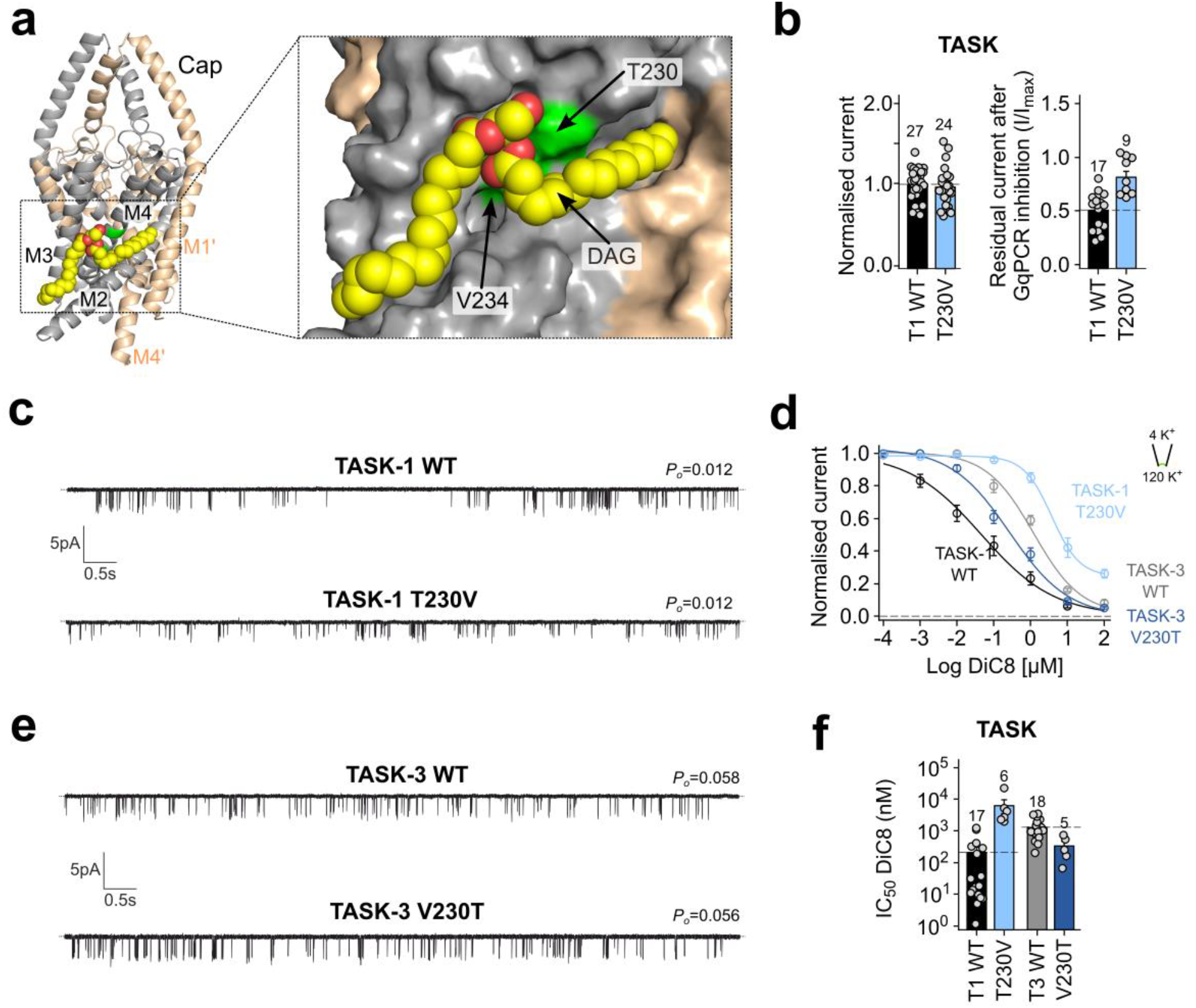
A residue on M4 defines the GqPCR and DAG sensitivity of TASK channels. **a**. Cartoon representation of the structure of TASK-1 with DAG bound. The two subunits are coloured grey and wheat with DAG shown in yellow as vdW spheres with cpk colouring. DAG binds within a groove between M2, M3 and M4 of the same subunit. The expanded panel on the right shows the groove in more detail with TASK-1 in surface representation. The key residues on M4 which influence DAG inhibition (T230 and V234) are shown in green. **b**. The T230V mutation in TASK-1 reduces GqPCR sensitivity but does not affect whole cell currents. **c**. The TASK-1 T230V mutation does not affect single channel *P*_*o*_ (mean values of *P*_*o*_ for WT and T230V mutant channels were 0.015±0.003 (n=8) and 0.012±0.003 (n=8), respectively). **d**. Dose-response curves for WT and mutant TASK-1 and TASK-3 channels showing effect of mutations on DiC8 sensitivity. **e**. The V230T mutation in TASK-3 does not affect single channel *P*_*o*_ (mean values of *P*_*o*_ for WT and V230T mutant channels were 0.058±0.008 (n=8) and 0.056±0.012 (n=8), respectively **f**. IC_50_ values from data in panel d showing the opposing effect of mutations at residue 230 on DiC8 sensitivity of TASK-1 (T1) and TASK-3 (T3).

Our results above demonstrate that any mutation which affects channel *P*_*o*_, can influence GqPCR inhibition irrespective of whether it may form part of the binding site or not. Therefore, direct effects of a mutation on DAG binding can only be inferred from a mutation which alters GqPCR sensitivity without significant changes in single channel *P*_*o*_. However, to simplify the process, an initial screen was first made for the effect of mutations within this groove on macroscopic whole-cell currents and GqPCR sensitivity before then examining their single channel *P*_*o*_ (**Supplementary Figure S1b**).

From this initial screen, several mutations were identified which reduced GqPCR sensitivity, but many either also increased the whole-cell currents, or were subsequently found to increase single channel *P*_*o*_. However, both the T230V and V234L mutations within the proposed site produced a marked reduction in GqPCR sensitivity, but whereas the V234L mutation increased channel *P*_*o*_, albeit only 2-fold (**Supplementary Figure S1c**), no increase in *P*_*o*_ was observed for the T230V mutation compared to WT TASK-1 (**Figure 5b-c**). Both the T230V and V234L mutations also retained their sensitivity to inhibition by external H^+^ suggesting that their principal gating processes remain otherwise intact (**Supplementary Figure S1d**)

### A binding site that defines the GqPCR sensitivity of TASK-1 and TASK-3

Due to its clear effect on GqPCR sensitivity without any increase in *P*_*o*_, the T230V mutation was therefore chosen for further investigation in macroscopic excised patch recordings and found to result in a dramatically reduced DAG sensitivity with the T230V mutant channel IC_50_ shifted ~100-fold compared to WT TASK-1 (**Figure 5d**). Furthermore, even though the V234L mutant produced a small increase in *P*_*o*_ it also resulted in a marked (>100-fold) decrease in DAG sensitivity (**Supplementary Figure S1e**). This therefore strongly supports a role for this groove in the interaction with DAG and its inhibitory effect, with a key role for residue T230 in particular.

Interestingly, TASK-1 and TASK-3 exhibit around 60-70 % amino acid sequence identity, especially within the core transmembrane regions and in M2 their sequence only differs by three amino acids, one of which is T230. In TASK-3 this residue is a valine (V230) and this channel has a reduced GqPCR sensitivity compared to TASK-1. We therefore examined the effect of mutating V230 on the GqPCR sensitivity of TASK-3 and found that the reverse (V230T) mutation increased GqPCR sensitivity with a concomitant increase in DAG sensitivity (**Figure 5d**). Importantly, the V230T mutation did not affect single channel *P*_*o*_ (**Figure 5e**). These results therefore strongly support a key role for this residue in the interaction of TASK channels with DAG and in defining their relative sensitivity to GqPCR-mediated inhibition.

## Discussion

In this study, we have identified a site within the membrane-facing transmembrane domains of TASK K2P channels that is critical for GPCR-mediated regulation of channel activity. This site interacts with DAG, a product of PIP_2_ hydrolysis by PLC, to mediate the effect of Gαq-coupled receptors. Our results provide a model whereby DAG interaction with this site results in a state-dependent inhibition, with DAG preferentially destabilising the open state of the channel to reduce channel *P*_*o*_. We also show that this mechanism is the primary reason why all of the known disease-causing GoF mutations produce channels with a markedly reduced GPCR sensitivity. This not only exacerbates the enhanced current levels produced by these mutations, but perhaps more importantly, also uncouples them from regulation by GqPCR-mediated signalling pathways. Our results therefore provide a greater insight into the mechanisms that control of TASK K2P channel gating, as well identifying a common regulatory defect shared by this GoF class of pathogenic mutations.

Previous studies have shown that DAG directly inhibits both TASK-1 and TASK-3 channels (*14, 15*), but the precise molecular mechanisms and site of action remained unclear. Likewise, it was not fully understood why all of the pathogenic GoF mutations, or indeed any mechanism that activates either TASK-1 or TASK-3 results in a dramatic reduction in GqPCR-mediated inhibition. Such inhibition is critically important for neuronal activity in the control of sleep, chemosensation and ventilation, as well as for vasomotor tone and aldosterone secretion. The regulation of TASK channel activity is also proposed as a potential mechanism for the treatment of atrial fibrillation and sleep apnea. Greater insight into these mechanisms of regulation therefore has the potential to improve exploitation of these channels as therapeutic targets.

We demonstrate a clear inverse correlation between single channel *P*_*o*_ and GPCR sensitivity which is also reflected in corresponding reductions in DAG sensitivity. Furthermore, we show that dynamic modulation of TASK-1 channel activity by either pH_e_ or by an agonist such as NS3623 also modulates GPCR inhibition via changes in DAG sensitivity thus supporting a state-dependent mechanism of action of DAG on overall TASK channel activity (**Figure 6**).

**Figure 6.**
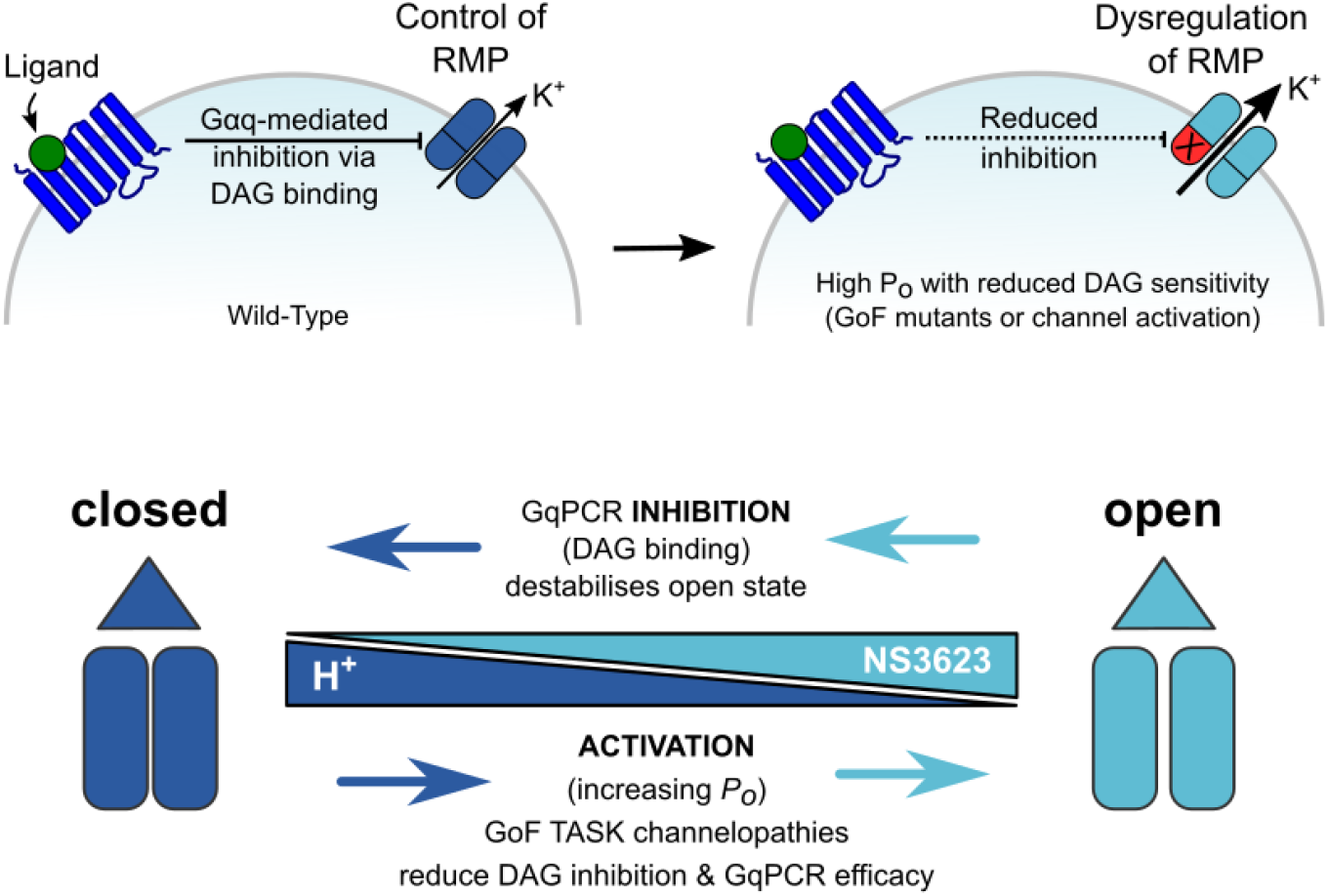
A model for GqPCR regulation of TASK channels and their dysregulation in disease. (Left) Cartoon describing GqPCR inhibition of TASK channels: GqPCR activation results in an increase in DAG levels (via hydrolysis of PIP_2_) and direct inhibition of TASK activity. Inhibition of TASK K^+^ channel activity results in depolarisation of the resting membrane potential (RMP) thus linking GqPCR activity to cellular electrical activity. DAG exhibits state-dependent regulation so GoF mutations that increase *P*_*o*_ reduce DAG inhibition resulting in increased TASK channel activity and inability to control the RMP due to uncoupling from GqPCR pathways. (Right) Dysregulation arises via state-dependent inhibition. DAG preferentially destabilises the open states of TASK channels so any mutant that increase *P*_*o*_ or mechanism that increases activity (*e*.*g*. change in pH_e_ or drugs such as NS3623) will reduce DAG-mediated inhibition and GqPCR efficacy.

Our single channel studies show that direct application of DiC8, a short chain DAG analogue preferentially destabilises the open state of the channel. Ideally, a detailed kinetic model of single channel gating would be developed to explore these effects in greater detail, but the low intrinsic *P*_*o*_ of TASK-1 means such analysis is not easily achievable. Nevertheless, the results clearly reveal a state-dependent mechanism of action that explains why channel activation, either resulting from pathogenic mutations or from dynamic channel modulation itself reduces TASK channel sensitivity to GPCR-mediated inhibition.

This state-dependent mechanism also helps to explain the pathogenic effect of activatory (*i*.*e*., GoF) mutations in both TASK-1 and TASK-3 channelopathies. In all these disease-causing variants, there is a marked increase in channel *P*_*o*_, and in the majority of cases increased whole-cell currents as well. However, in some variants e.g., L241F within the X-gate of TASK-1, there is little effect on the magnitude of whole-cell currents due to an associated reduction in channel trafficking to the plasma membrane (*15, 22*) indicating that it may be the lack of GqPCR regulation itself that is the principal pathogenic effect.

Our results also provide insight into the binding site for DAG on TASK channels and for the differences in the GqPCR sensitivity of TASK-1 and TASK-3. The MD simulations reveal DAG interaction with an external membrane-facing groove formed by M2, M3 and M4 of the same subunit of TASK-1, including a critically important interaction with T230 on M4. Interestingly, threonine residues are not uncommon within transmembrane helices due to their ability to influence helical bends (*36*). This site is also located just above the X-gate on M4 within the membrane and is therefore well placed to influence the relative stability of the open vs closed state of the channel. Indeed, it has been shown that DAG is highly mobile within the membrane and even able to accumulate between leaflets (*35*) thus permitting ready access of DAG to this site after PIP_2_ hydrolysis.

No structure yet exists for an open TASK channel and so our structural modelling only examines the closed state, but the X-gate resides below T230 on M4 and only relatively small movements or rotations of M4 at this site are likely to be required to open the X-gate. Thus, the interaction of the DAG headgroup with T230 combined with other non-polar interactions may therefore be sufficient to destabilise the open state. Importantly, T230 in TASK-1 is also one of the few amino acids that is different in TASK-3 within this region, and we show that exchanging this single residue has opposing effects on GqPCR sensitivity without any changes in *P*_*o*_ thus explaining why TASK-3 channels are intrinsically less sensitive to GqPCR inhibition.

In conclusion, our findings indicate that binding of DAG to an external-facing groove within the TM helices, to preferentially destabilise the open state, underlies the inhibition of TASK channels via GqPCR signalling. This state-dependent mechanism explains the common defect found in many TASK channelopathies and not only advances our understanding of GPCR regulation of TASK channels but also the pathogenic effect of TASK channelopathies that result from channel dysregulation.

## Acknowledgments & Funding Information

This work was directly supported by grants from the Biotechnology and Biological Sciences Research Council and Medical Research Council to S.J.T and P.C.B. (BB/T002018/1, BB/S008608/1, BB/S001247/1 and MR/W017741/1). It was also supported by the Wellcome Trust as part of the OXION Initiative in Ion Channels and Membrane Transport in Health and Disease (WT084655MA and 102161/B/13/Z). Further grants from the Deutsche Forschungsgemeinschaft supported the work of M.S. (SCHE 2112/1-2) and T.B (BA 1793/6-2) as part of the Research Unit FOR2518, *DynIon*. It was also supported by the Leibniz Collaborative Excellence Programme (K622/2024) to T.B and M.S. D.S. and K.C. are funded by the UKRI-BBSRC Interdisciplinary Bioscience Doctoral Training Partnership (BB/M011224/1).

## Author contributions

S.J.T., T.H.J-T, P.P. and M.S. conceived/designed the principal elements of the study. All authors (T.H.J-T., P.P., D.S., K.C., P.C.B., T.B., M.S., and S.J.T.) generated, analyzed or interpreted data, or generated materials. T.H.J-T, S.J.T, and M.S. drafted the manuscript and all authors contributed to the final version.

## Supplementary Information

**Supplementary Table S1.**
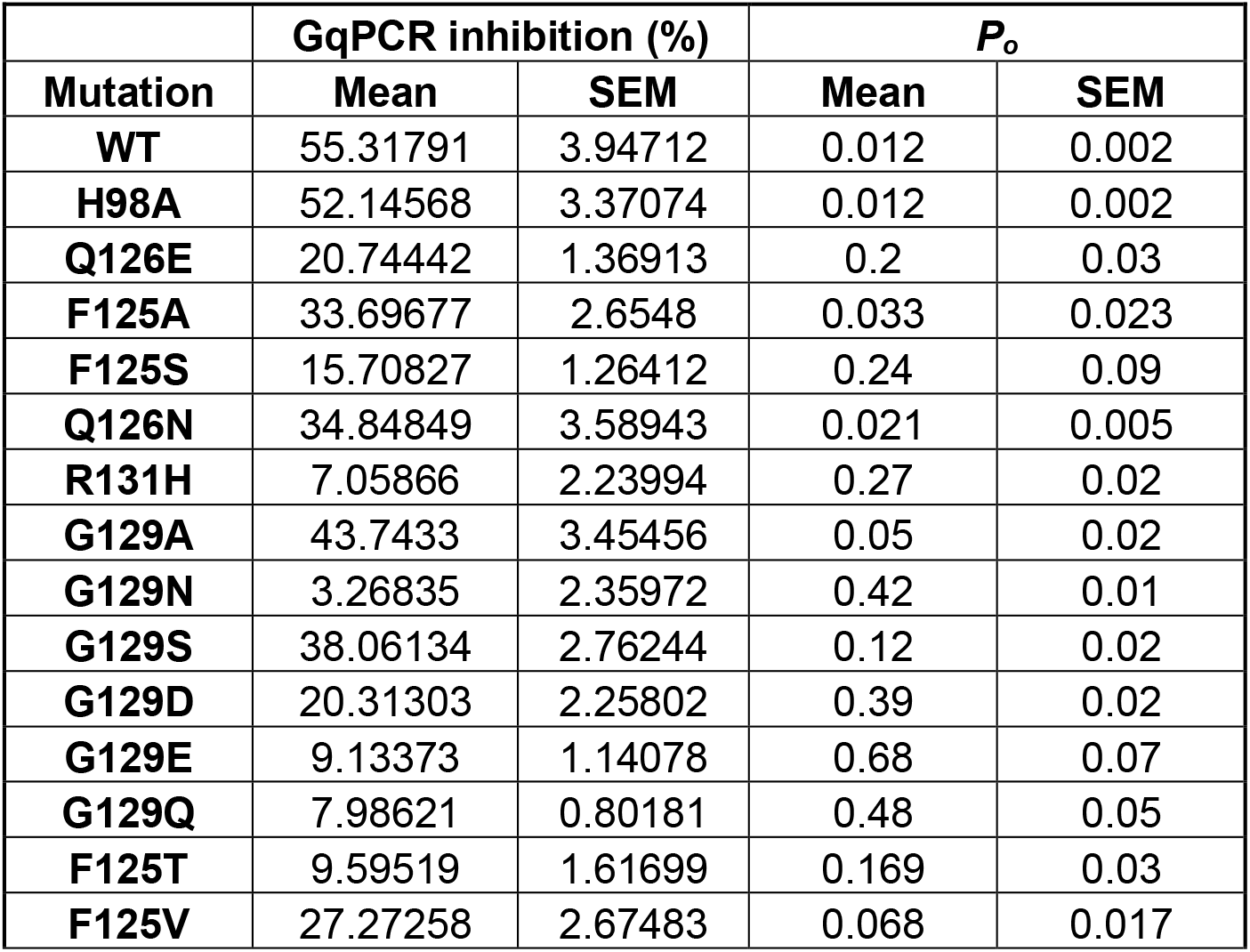
Values for GqPCR inhibition and single channel *Po* for the mutations shown in Figure 1d.

| | GqPCR inhibition (%) | | $P_o$ | |
| --- | --- | --- | --- | --- |
| Mutation | Mean | SEM | Mean | SEM |
| WT | 55.31791 | 3.94712 | 0.012 | 0.002 |
| H98A | 52.14568 | 3.37074 | 0.012 | 0.002 |
| Q126E | 20.74442 | 1.36913 | 0.2 | 0.03 |
| F125A | 33.69677 | 2.6548 | 0.033 | 0.023 |
| F125S | 15.70827 | 1.26412 | 0.24 | 0.09 |
| Q126N | 34.84849 | 3.58943 | 0.021 | 0.005 |
| R131H | 7.05866 | 2.23994 | 0.27 | 0.02 |
| G129A | 43.7433 | 3.45456 | 0.05 | 0.02 |
| G129N | 3.26835 | 2.35972 | 0.42 | 0.01 |
| G129S | 38.06134 | 2.76244 | 0.12 | 0.02 |
| G129D | 20.31303 | 2.25802 | 0.39 | 0.02 |
| G129E | 9.13373 | 1.14078 | 0.68 | 0.07 |
| G129Q | 7.98621 | 0.80181 | 0.48 | 0.05 |
| F125T | 9.59519 | 1.61699 | 0.169 | 0.03 |
| F125V | 27.27258 | 2.67483 | 0.068 | 0.017 |

**Supplementary Table S2.** Mean lifetimes (τ) and relative areas (*a*) of open states, closed states and burst durations of the probability density functions fitted to the data in Figure 4.

| | $\tau$ (ms) (control) | $\tau$ (ms) 1 μM DiC8 | $a$ (control) | $a$ (1 μM DiC8) |
| --- | --- | --- | --- | --- |
| Open states | 0.65 | 0.35 | 0.24 | 0.88 |
|  | 1.29 | 0.63 | 0.76 | 0.12 |
| Closed states | 0.15 | 0.04 | 0.75 | 0.12 |
|  | 0.26 | 0.15 | 0.18 | 0.18 |
|  | 1.23 | 1.1 | 0.02 | 0.04 |
|  | 10.5 | 23 | 0.008 | 0.22 |
|  | 41 | 56 | 0.036 | 0.44 |
|  | 620 | 330 | 0.0003 | 0.003 |
| Bursts | 0.10 | 0.41 | 0.04 | 0.67 |
|  | 1.6 | 0.86 | 0.35 | 0.23 |
|  | 24.5 | - | 0.61 | - |

**Supplementary Figure S1.**
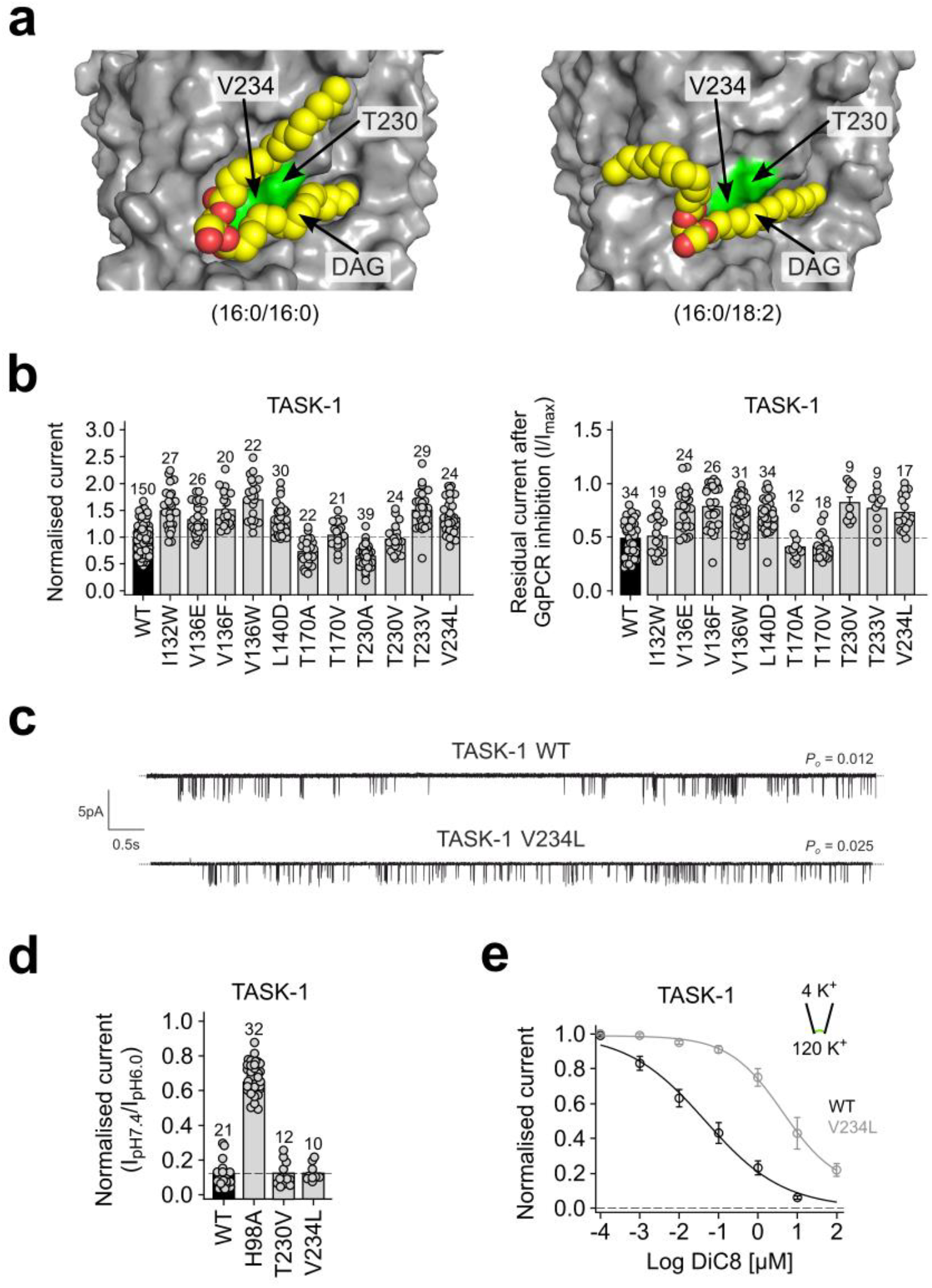
Molecular determinants of DAG sensitivity of TASK channels. **a**. Less frequent poses also found for DAG in the groove between M2, M3 & M4 (see also Figure 5a). Residues T230 and V234 are shown in green. **b**. Normalised whole cell current values and relative GqPCR sensitivity for mutations within this groove. **c**. Single channel recordings for the V234L mutation in TASK-1. This mutation increases *P*_*o*_ ~2-fold. **d**. Both T230V and V234L within this binding site retain their sensitivity to inhibition by external pH. The H98A mutation of the pH sensor is shown as a control. **e**. Despite its small increase in *P*_*o*_, the V234L mutation causes a marked reduction in DiC8 sensitivity as measured in giant excised patches.

